# Variations in neuromodulatory chemical signatures define compartments in macaque cortex

**DOI:** 10.1101/272849

**Authors:** Nicholas J. Ward, Wolf Zinke, Jennifer J. Coppola, Anita A. Disney

## Abstract

Subcortical neuromodulatory systems exhibit widespread projections that influence the output of cortical circuits. These modulatory systems have been portrayed as providing global signals to the entirety of cortex based on their widespread innervation. This innervation is not necessarily predictive of the levels of the neuromodulatory molecules that actually provide these signals to cortex. In the present study, we examine tissue concentrations of dopamine, noradrenaline, and serotonin to see how they locally vary across multiple areas of the macaque cortex. Our results indicate that different cortical areas exhibit varying levels of dopamine, noradrenaline, and serotonin. Using cluster analysis, we examine how similar cortical regions are to each other, finding that similarities in neurochemical content are shared by areas that exhibit similar functionality. Altogether, these findings demonstrate that neurochemical signatures vary across cortical regions and help define unique, local neuromodulatory signaling compartments.

## Introduction

In examining how subcortical neuromodulatory nuclei influence cortex, early interpretations position these modulatory systems as general influencers or conveyors of global signals (Saper 1987). Such roles in cortical signaling have been ascribed to the cholinergic (Phillis 1968), dopaminergic (Schultz 2002), noradrenergic (Freedman et al. 1975, Jones and Moore 1977, Gatter and Powell 1977), and serotonergic (Wilson and Molliver 1991) systems. Assertions of global neuromodulatory signaling are often based on observation of diffuse axonal innervation of the cortex by these subcortical nuclei; however, innervation density and pattern do not describe the modulatory signal itself. The modulatory signal is instead represented by the moment-to-moment extracellular concentrations of the molecules those axons release: acetylcholine, dopamine, noradrenaline, and serotonin. It is not known whether extracellular concentration can be predicted based on innervation density, but there are data to support why this might not be the case (reviewed by Coppola et al. 2016). In the end, the ability to assess hypotheses related to the global vs. local nature of signaling requires direct measurement of these neurochemicals in cortex.

Efforts to understand cortical neuromodulation in non-human primates and humans are currently dominated by approaches that detail the circuit elements surrounding neuromodulatory signals. In histological studies, influences of neuromodulatory systems are extrapolated from descriptions of axonal innervation by subcortical modulatory nuclei and receptor expression by the receiving cortical circuit. Electrophysiological studies can describe activity of neurons in the neuromodulatory nuclei and the response of the cortical circuits into which these nuclei send axons. Often absent from both types of study is a description of the concentration dynamics of the signaling molecules themselves. Without information about the molecules that transfer the signal between innervating axon and receiving circuit, the anatomy of innervation patterns and receptor expression is not sufficient for gaining a full understanding of modulatory signaling. Neuromodulatory systems signal partly through volume transmission, wherein molecules are released from varicosities not apposed to a receptive surface (Fuxe and Agnati 1991). These molecules instead diffuse over some distance in the extracellular space, activating nearby receptors (similar to synaptically-released molecules that “spill over” outside of the synapse). In this signaling mode, the relationship between signal sent and signal received becomes vulnerable to extrasynaptic influences and can be readily weakened (Coppola et al. 2016). As such, it is necessary to measure the concentration dynamics of the signaling molecules themselves. Construction of *in vivo* dynamical studies is informed by non-dynamical, or steady-state, measurements of modulator concentration in postmortem frozen tissue. Compared with histological and electrophysiological studies of neuromodulatory systems in non-human primates, fewer studies detail steady-state measures of multiple modulators across cortex (Brown et al. 1979, Pifl et al. 1991).

This paper addresses the need for steady-state measures of modulators across cortical regions. Using unfixed frozen tissue from macaque cortex, we quantified concentrations of monoaminergic neuromodulatory molecules in several regions. We assert that these measured concentrations, taken together, represent a neuromodulatory chemical signature that differs across cortex. These neurochemical signatures, when combined with known local circuit features, can be used to understand the cortex as being composed of modulatory compartments (Coppola et al. 2016).

## Materials and Methods

### Animals

Four adult (9.25-11.5 years old) male rhesus macaques (*Macaca mulatta*) were examined for cortical concentrations of biogenic monoamines. One of the four (macaque 16) had been a subject in prior experimental studies, the remaining three were naïve animals (macaques 13-15). Macaque 16 was previously a subject both in behavioral experiments detailed by Dylla et al. (2013) and in physiological studies. Electrodes in these single-unit recording studies targeted the inferior colliculus on the right hemisphere and the cochlear nucleus on the left hemisphere. Electrode tracts passed through parietal cortex and potentially extrastriate visual cortical areas (for inferior colliculus recordings) and lateral visual cortex, possibly including striate cortex (for cochlear nucleus recordings). Experimental procedures were approved by the Vanderbilt Institutional Animal Care and Use Committee and followed the guidelines published by the National Institutes of Health.

### Tissue sample acquisition

Anesthesia was induced with ketamine (10 mg/kg) and midazolam (0.05 mg/kg) and supplemented (post-induction) with ketamine (4 mg/kg) and dexmedetomidine (5 μg/kg). Animals were then euthanized by sodium pentobarbital overdose (120 mg/kg), and when areflexive, a craniectomy was performed. The brain was removed from the skull without exsanguination or fixation, to allow for fresh tissue analysis. An unrelated study constrained regions of the brain available for our analyses to portions of the left hemispheres of macaques 13 and 16 and the right hemispheres of macaques 14 and 15. These hemispheres were first separated into two or three larger blocks: either frontal/middle and caudal blocks or frontal, middle (from near the temporal pole to posterior at the level of the cochlear nucleus), and caudal blocks. From these blocks, coronal tissue slabs were dissected to include specific regions of cortex. Slab collection from macaque 16 avoided tissue potentially damaged during electrophysiological recording. Following dissection, tissue slabs were immediately submerged in plastic trays of optimal cutting temperature (OCT) compound (Tissue-Tek, Sakura Finetek, Torrance, CA, USA) and frozen on dry ice for storage at −80°C prior to assay. All four acquisition procedures each lasted 30-40 minutes from beginning of craniectomy to freezing of the last tissue slab. Tissue punches were collected from frozen cortical tissue slabs in preparation for biogenic monoamine analysis.

### Defining regions of cortex

Regions of sampled tissue are represented in **Figure 1**. Samples collected from each animal are detailed in **Table 1**. Motor (M1) and somatosensory (S1) cortex tissue was collected by making an incision along the depth of the central sulcus. “M1” blocks comprised tissue anterior to this incision and posterior to the precentral dimple. “S1” blocks comprised tissue between the central sulcus and the postcentral dimple. Two slabs containing the central sulcus and tissue immediately surrounding it were collected from the dorsomedial (SMd) and ventrolateral (SMv) portions of the sulcus. Anterior to the slabs that sampled these sensorimotor areas (M1, S1, SMd, SMv), tissue containing premotor areas (PMA) was collected posterior to the genu of the arcuate sulcus. Posterior parietal cortex was broadly divided into two slabs, separated by the intraparietal sulcus. Dorsal and ventral posterior parietal cortex (dPPC and vPPC, respectively) tissue was dissected using slabs containing the superior and inferior parietal lobules, respectively. Primary and secondary visual cortex (V1 and V2, respectively) tissue was collected from slabs containing the calcarine sulcus as a reference point. Further details regarding these samples are found in **Table 1**. Extrastriate visual area 3 (V3) tissue was collected from both dorsal (V3d) and ventral (V3v) regions. V3d tissue was collected from slabs containing the inferior bank of the lunate sulcus. V3v tissue was collected from slabs containing the ventral surface of cortex near the caudal end of the occipitotemporal sulcus and laterally to the inferior occipital sulcus.

**Figure 1.**
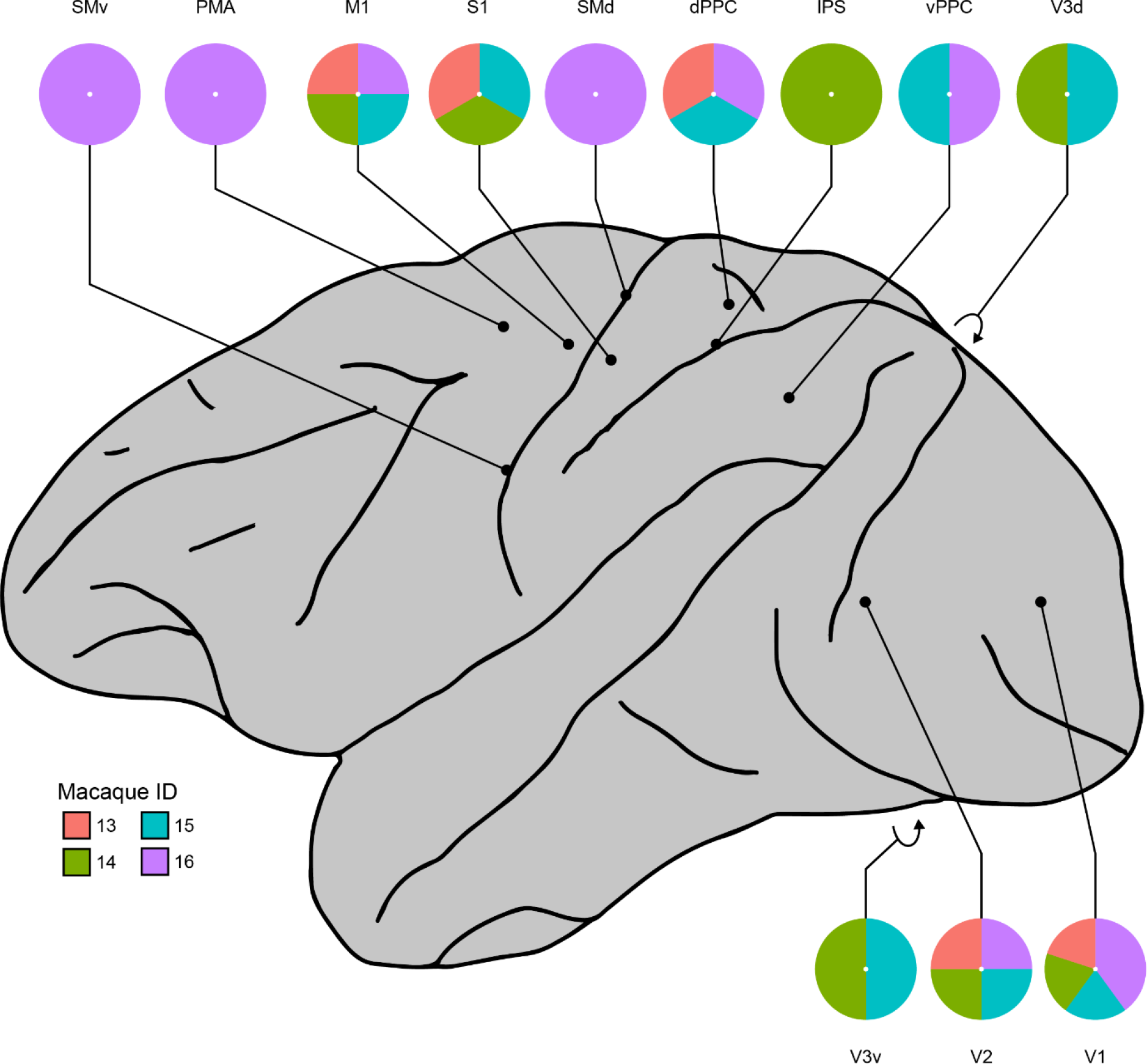
Cortical areas sampled for analysis of monoamine concentrations. For each area in cortex, colors in pinwheel represent macaques sampled for that area. Map of lateral cortical surface adapted from Romanski et al. (2005). V1, primary visual cortex; V2, secondary visual cortex; V3v, extrastriate visual area 3 (ventral); V3d, extrastriate visual area 3 (dorsal); vPPC, ventral posterior parietal cortex; dPPC, dorsal posterior parietal cortex; IPS, intraparietal sulcus; SMv, sensorimotor cortex (ventral); SMd, sensorimotor cortex (dorsal); S1, primary somatosensory cortex; M1, primary motor cortex; PMA, premotor areas.

**Table 1.** Tissue samples collected from each cortical area.

| Cortical area | Macaque ID | Tissue slab contents | Samples collected |
| --- | --- | --- | --- |
| Premotor areas (PMA) | 16 |  | 3 |
| Primary motor cortex (M1) | 13 |  | 3 |
|  | 14 |  | 3 |
|  | 15 |  | 3 |
|  | 16 | Motor cortex and its caudal border | 3 |
| Sensorimotor cortex, dorsomedial (SMd) | 16 | Ranging from middle to dorsal-most extent of central sulcus | 3 |
| Sensorimotor cortex, ventrolateral (SMv) | 16 | Ranging from middle to lateral-most extent of central sulcus, including some rostral portions of posterior parietal cortex | 3 |
| Primary somatosensory cortex (S1) | 13 |  | 3 |
|  | 14 |  | 3 |
|  | 15 |  | 3 |
| Intraparietal sulcus (IPS) | 14 | Intraparietal sulcus and surrounding tissue | 3 |
| Posterior parietal cortex, dorsal (PPCd) | 13 | May contain some of Area 2 and half of the dorsal posterior parietal cortex | 3 |
|  | 15 |  | 3 |
| Posterior parietal cortex, ventral (PPCv) | 15 |  | 3 |
| Primary visual cortex (V1) | 13 | Foveal and perifoveal V1 | 3 |
|  | 14 | Foveal and perifoveal V1 | 3 |
|  | 15 | Mostly perifoveal V1 | 3 |
|  | 16 | Calcarine sulcus |  |
|  | 16 | Only V1, caudal pole | 3 |
| Visual area 2 (V2) | 13 | Half of the calcarine sulcus, V2, and potentially some V3 | 3 |
|  | 14 | V1/V2 border, and most of V2 | 3 |
|  | 15 | V1/V2 border, V2, and some dorsal V3 | 3 |
|  | 16 | V1 and V2 | 3 |
| Visual area 3, dorsal (V3d) | 14 | Dorsal V3 and some dorsal posterior parietal cortex | 3 |
|  | 15 |  | 3 |
|  | 16 |  | 3 |
|  | 15 |  | 3 |
| Visual area 3,<br>ventral (V3v) | 16 |  | 3 |

### Tissue punch extraction

Tissue punches were collected by penetrating the tissue slab with hollow hypodermic tubing (3 mm inner diameter). Using a scalpel blade, OCT was pared away from either end of the sample and obvious white matter was removed such that each sample was mostly representative of gray matter tissue. Prior to collection, materials (scalpel, tubing, etc.) were sterilized to reduce contamination or degradation of samples. After sampling, both the collected tissue punches and tissue slab were returned to dry ice for temporary storage during sampling, and then subsequently to a −80°C freezer.

Cortical tissue punches were homogenized using a tissue dismembrator in 100-750 μl of a solution containing 0.1M trichloroacetic acid (TCA), 10^−2^ M sodium acetate, 10^−4^ M EDTA, and 10.5% methanol (pH 3.8). Ten microliters of homogenate were used for protein assay. Samples were spun in a microcentrifuge at 10,000 g for 20 minutes. The supernatant was removed for biogenic monoamine analysis.

### Biogenic amine analysis using high performance liquid chromatography with electrochemical detection (HPLC-ECD)

Levels of biogenic amines were determined using an Antec Decade II (oxidation: 0.65) electrochemical detector operated at 33°C. Twenty microliter samples of the supernatant were injected using a Waters 2707 autosampler onto a Phenomenex Kintex C18 HPLC column (100 × 4.60 mm, 2.6 μm). Biogenic amines were eluted with a mobile phase consisting of 89.5% 0.1M TCA, 10^−2^ M sodium acetate, 10^−4^ M EDTA and 10.5% methanol (pH 3.8). Solvent was delivered at 0.6 ml/min using a Waters 515 HPLC pump. Using this HPLC solvent, the biogenic amines elute in the following order: noradrenaline, adrenaline, DOPAC, dopamine, 5-HIAA, HVA, 5-HT, and 3-MT. HPLC control and data acquisition are managed by Empower software. Isoproterenol (5 ng/ml) was included in the homogenization buffer as a standard to quantify the biogenic amines.

### Biogenic amine analysis using liquid chromatography/mass spectrometry (LC/MS)

Levels of biogenic amines were determined using LC/MS following derivatization of analytes with benzoyl chloride (BZC). Five microliters of tissue extract was added to a 1.5 ml microcentrifuge tube containing 20 μl acetonitrile. Ten microliters each of 500 mM aqueous NaCO_3_ and 2% BZC in acetonitrile was added to each tube. After two minutes, the reaction was stopped by the addition of 20 μl internal standard solution (in 20% acetonitrile containing 3% sulfuric acid) and 40 μl water.

LC was performed on a 2.0 × 50 mm, 1.7 μm particle Acquity BEH C18 column (Waters Corporation, Milford, MA, USA) using a Waters Acquity UPLC. Mobile phase A was 15% aqueous formic acid and mobile phase B was acetonitrile. Samples were separated by a gradient of 98–5% of mobile phase A over 11 min at a flow rate of 600 μl/min prior to delivery to a SCIEX 6500+ QTrap mass spectrometer.

### Protein assay

Protein concentration was determined by bicinchoninic acid (BCA) Protein Assay Kit (Thermo Scientific). In individual wells of a 96-well plate, 10 μl of sample tissue homogenate and 200 μl of BCA working reagent were added. The plate was incubated at room temperature for two hours for color development. A bovine serum albumin standard curve was run at the same time. Absorbance was measured using a POLARstar Omega plate reader (BMG LABTECH Company).

### Analysis and Statistics

All statistical analyses were conducted using R version 3.4.3 (R Core Team 2017). Graphs were plotted in R using ggplot2 (Wickham 2009), scatterplot3d (Ligges and Mächler 2003), cowplot (Wilke 2017), dendextend (Galili 2015) packages, and imported into Adobe Illustrator for stylistic adjustments. Correlations between HPLC and LC/MS measures for the same samples were assessed using Pearson’s product-moment correlation analyses. For plots where multiple data points were collapsed to visualize brain regions as single points, median values were used to represent center of mass (**Figures 4**, **5**).

Hierarchical clustering of brain regions was conducted using Ward’s hierarchical agglomerative clustering method (Ward 1963, Murtagh and Legendre 2014). Briefly, median monoamine concentration values for each cortical region were scaled, and dissimilarity between the regions was calculated as the Euclidean distance between pairs of these median values. At the beginning of the analysis, each cortical region was assigned to its own branch. With each iteration of the algorithm, the pair of branches with the lowest distance between each other was merged into a single branch until all branches were merged. An average silhouette analysis was conducted to estimate the number of clusters that best describe the data (Rousseeuw 1987). This analysis measures how similar an observation is to its own cluster as compared to other clusters.

## Results

We collected cortical tissue samples from frozen brains of adult, male macaques to examine how monoamine concentrations vary across cortex. Cortical regions sampled are indicated in **Figure 1**, and additional details regarding samples are noted in **Table 1**. We measured concentrations of dopamine, noradrenaline, and serotonin in all samples using both HPLC and LC/MS to ensure that our results are robust across measurement techniques and to facilitate comparison with prior studies (which have tended to use HPLC). Correlation analyses indicate a strong, positive, monotonic relationship between measurements made using both techniques (**Figure 2**). Only monoamine concentration data derived by LC/MS was used for subsequent analysis because a small number of samples analyzed by HPLC exhibited interference that prevented measurements.

**Figure 2.**
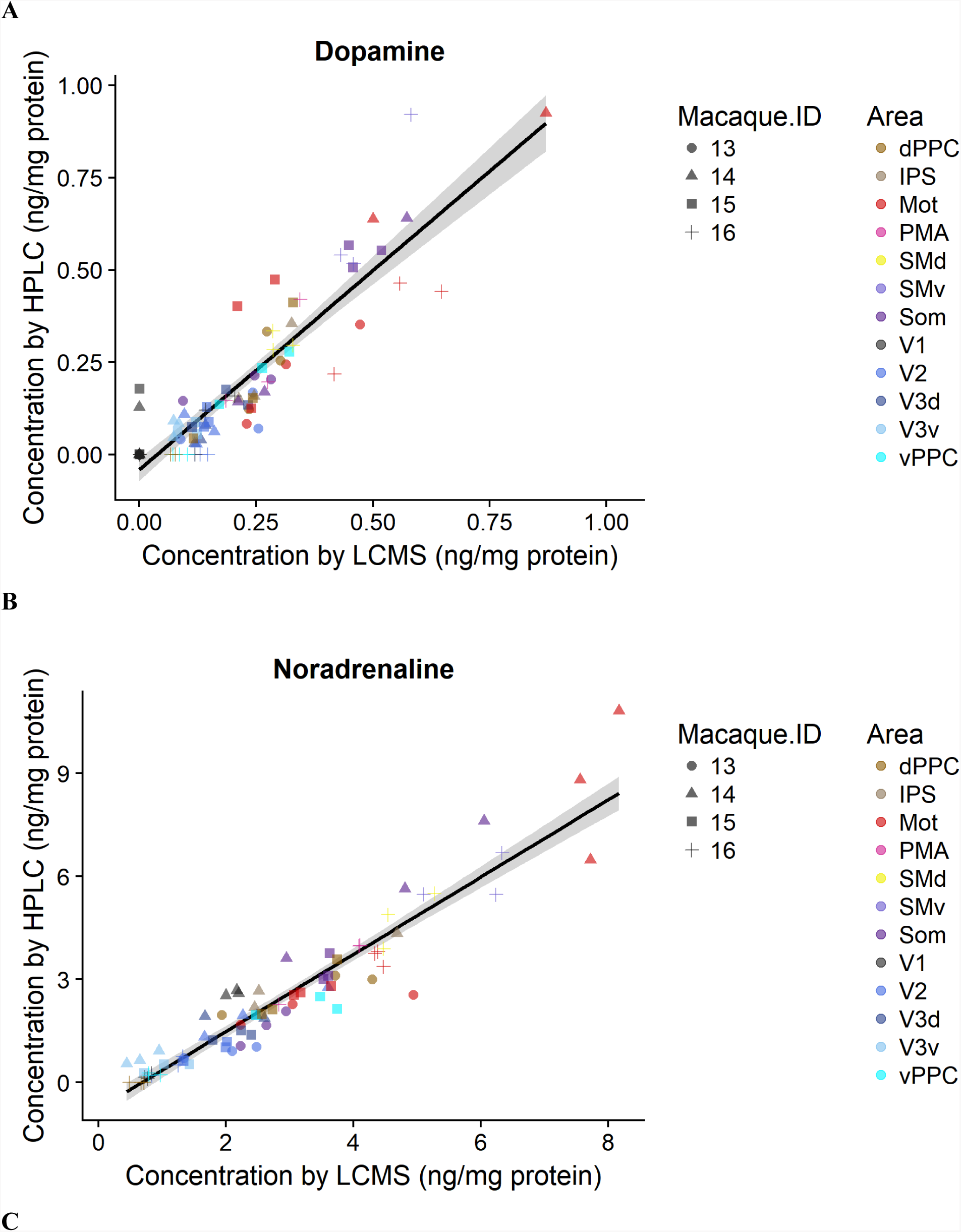
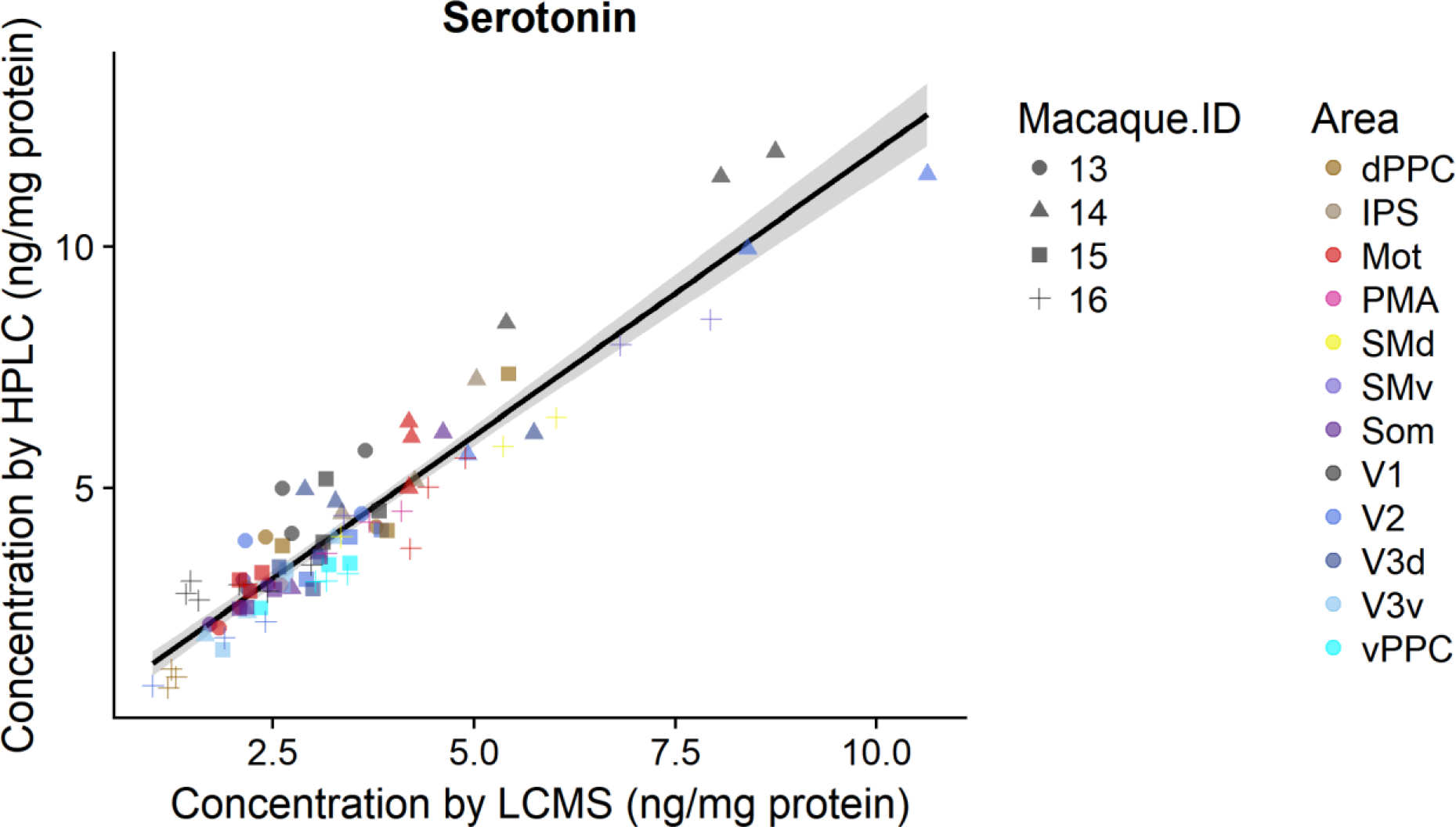
Correlation between HPLC and LCMS concentration measures for dopamine (**A**), noradrenaline (**B**), and serotonin (**C**).

LC/MS data were plotted in two-dimensional graphs of modulator pairs to examine how cortical regions vary in terms of neurochemical content (**Figure 3**). To aid interpretation of these relationships, all samples collected for a region were collapsed into a single, center-of-mass point using median values of all samples for that region (**Figure 4**). Values for all three monoamines were then plotted together to visualize the samples in a multi-modulator space (**Figure 5**).

**Figure 3.**
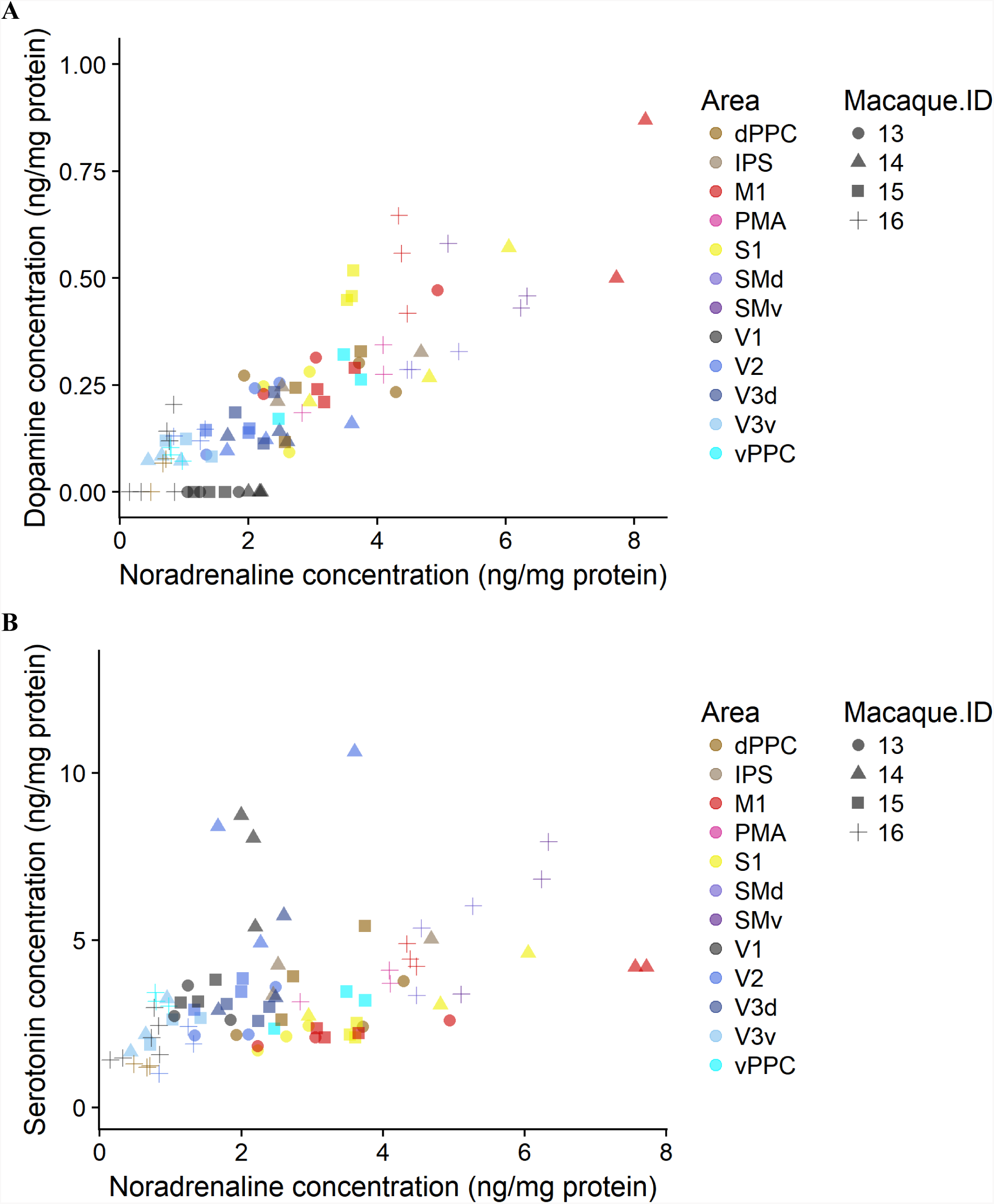
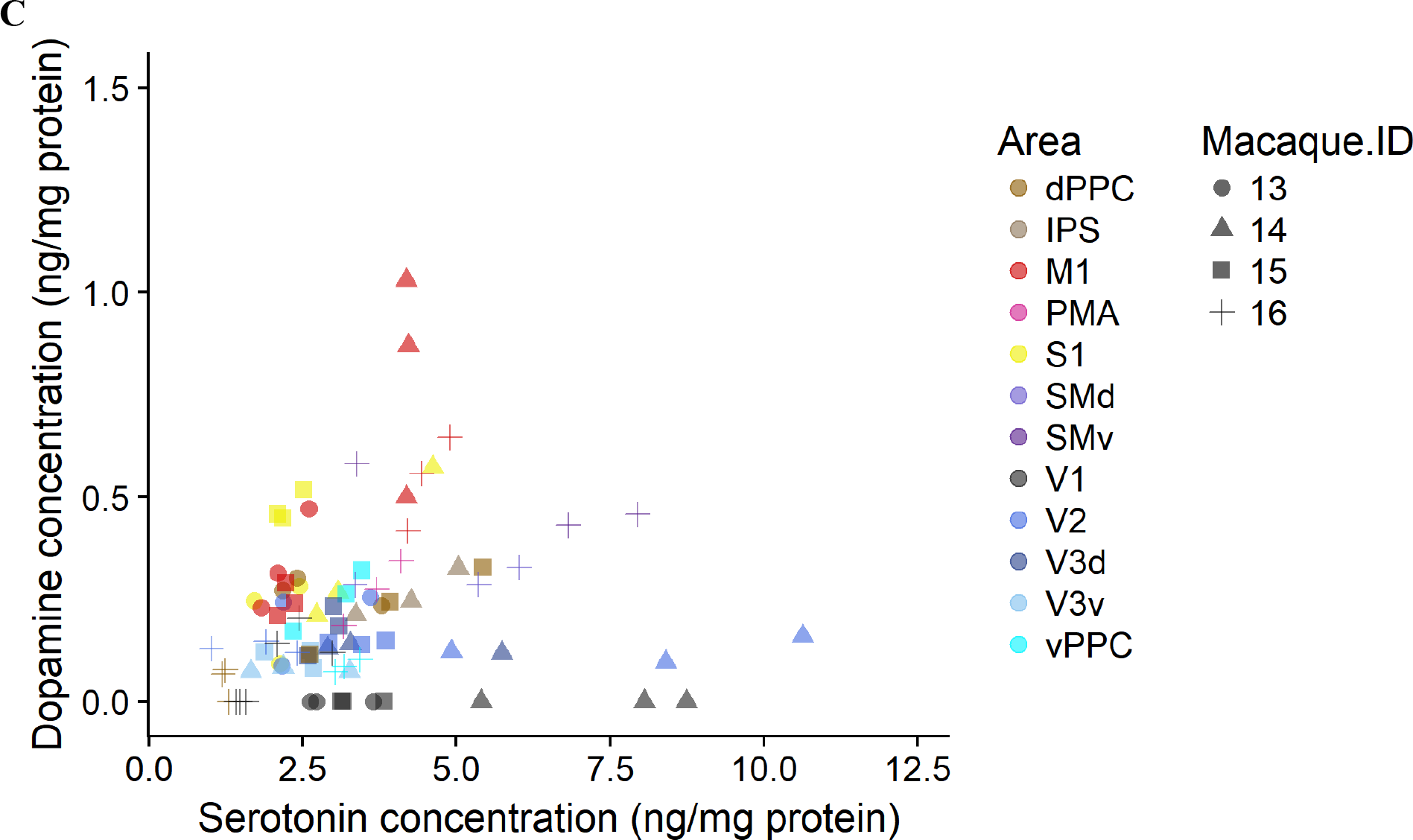
Plots showing monoamine concentration for each sample collected. As indicated in the legend, data points indicate the macaque (shape) and brain area (color) associated with the sample. Plots show comparisons of (**A**) dopamine and noradrenaline, (**B**) serotonin and noradrenaline, and (**C**) dopamine and serotonin.

To identify neurochemically similar cortical areas, we analyzed our monoamine concentration data by hierarchical clustering (**Figure 6**). Average silhouette analysis indicated that the data should be separated into two large clusters (**Figure 6**, boxes). One cluster was dominated by the visual areas (V1, V2, V3v, V3d) and also included PPCv. The other cluster was dominated by sensorimotor areas (M1, S1, SMv, SMd, PMA), and also included PPCd and IPS.

## Discussion

In the current study, we defined multidimensional neurochemical signatures in several areas of postmortem macaque cortex. To define these neurochemical signatures, we used LC/MS to measure monoamine concentrations in each cortical area. These concentrations were used to plot where each brain region exists within a three-dimensional representation of concentrations for dopamine, noradrenaline, and serotonin. We used cluster analysis to examine how similar cortical regions were to each other based on the distance from one region to another in this three-dimensional space. This cluster analysis suggested two main groupings: one that contained visual areas and PPCv, and another that contained sensorimotor areas as well as PPCd and IPS. These findings suggest that similarities in neurochemical signature are shared by areas that exhibit similar functionality. PPC and IPS samples split between these two main groupings, potentially as a result of the multimodal functions associated with these brain regions. To understand how these neurochemical signatures contribute to the existence of distinct neuromodulatory signaling environments, it is important to consider how these monoamines interact with local circuit elements. In this discussion, we will compare our results with other studies and examine how the neurochemical signatures we defined relate to features of local circuitry, including monoaminergic innervation patterns and expression of receptors across cortex. We will then expand upon how this modeling of neurochemical signatures can inform future approaches for *in vivo* study of brain state.

Cortical innervation density of neuromodulatory axons has been used as a proxy for understanding the neurochemical environment. The following three sections examine whether innervation serves as a useful proxy for understanding the neurochemical characteristics of a cortical region. **Table 2** summarizes this data across multiple studies, modulatory systems, and cortical regions. For ease of comparison across studies that use a variety of areal naming conventions, we have aligned cortical area nomenclature to those used in the present study.

**Table 2.** Summary of dopaminergic, noradrenergic, and serotonergic innervation density across multiple cortical areas.

| Dopaminergic innervation studies |  |  |  |  |  |  |  |  |  |
| --- | --- | --- | --- | --- | --- | --- | --- | --- | --- |
| Study | Somatosensory |  | Motor |  | Prefrontal |  | Parietal |  | Visual |
| Berger et al. (1988): macaque ([3H]DA) | S1 | < | M1 (medial>dorsal), premotor, supplementary motor | > | 46 | = | 7 | > | 18 > V1 |
| Gaspar et al. (1989): human (TH+) | S1 | < | M1, premotor | > | 9 | ? | 5 ? 40 | > | 18 > V1 |
| Levitt et al. (1984): macaque (SPG method, DA-like) | S1 | < | M1, premotor | ? | 9-14 | > | 5 ? 7 | ? | V1 ? 18 ? 19 |
| Lewis et al. (1987): macaque (TH+) | S1 | < | M1, premotor | > | 9 > 11, 12 > 16, 10 | ?/= | 7 | > | 18 > V1 |
| Noradrenergic innervation studies |  |  |  |  |  |  |  |  |  |
| Study | Somatosensory |  | Motor |  | Prefrontal |  | Parietal |  | Visual |
| Gaspar et al. (1989): human (DBH+) | S1 | = | M1 > premotor | > | 9 | = | 40 | = | 18 > V1 |
| Levitt et al. (1984): macaque (SPG method, NA-like) | S1 | > | M1 | ? | 9 ? 11 | > | 7 ? 19 | > | V1 ? 18 ? 19 |
| Morrison and Foote (1986): macaque and squirrel monkey (DBH+) | n.d. | ? | n.d. | ? | n.d. | ? | 5 = 7 | > | 18 > V1 |
| Lewis and Sesack (1997): macaque (DBH+) | S1 | > | M1 | > | 9 > 10 | ? | n.d. | ? | n.d. |
| Serotonergic innervation studies |  |  |  |  |  |  |  |  |  |
| Study | Somatosensory |  | Motor |  | Prefrontal |  | Parietal |  | Visual |
| Takeuchi and Sano (1983): macaque (5-HT+) | S1 | > | M1 | ? | n.d. | ? | n.d. | S1 < | 18 < V1 |
| Morrison and Foote (1986): macaque and squirrel monkey (5-HT+) | n.d. | ? | n.d. | ? | n.d. | ? | 5 = 7 | < | 18 < V1 |
| Berger et al. (1988): macaque ([3H]5-HT) | S1 | > | M1 | ? | 46 | ? | 5 | < | 18 < V1 |

### Comparisons between dopaminergic innervation and steady state measures

In a study of dopaminergic and serotonergic innervation of macaque cortex, all cortical areas examined by Berger et al. (1988) exhibited much sparser dopaminergic than serotonergic innervation, except M1 and anterior cingulate cortex. Our results reflect lower levels of dopamine than serotonin across the entire cortex, including M1 (**Figure 3A**). When comparing across areas, our M1 samples exhibit some of the highest levels of dopamine. Berger et al. (1988) additionally noted higher dopaminergic innervation of V2 than V1. Our results parallel this in terms of dopamine concentration (**Figure 4**).

**Figure 4.**
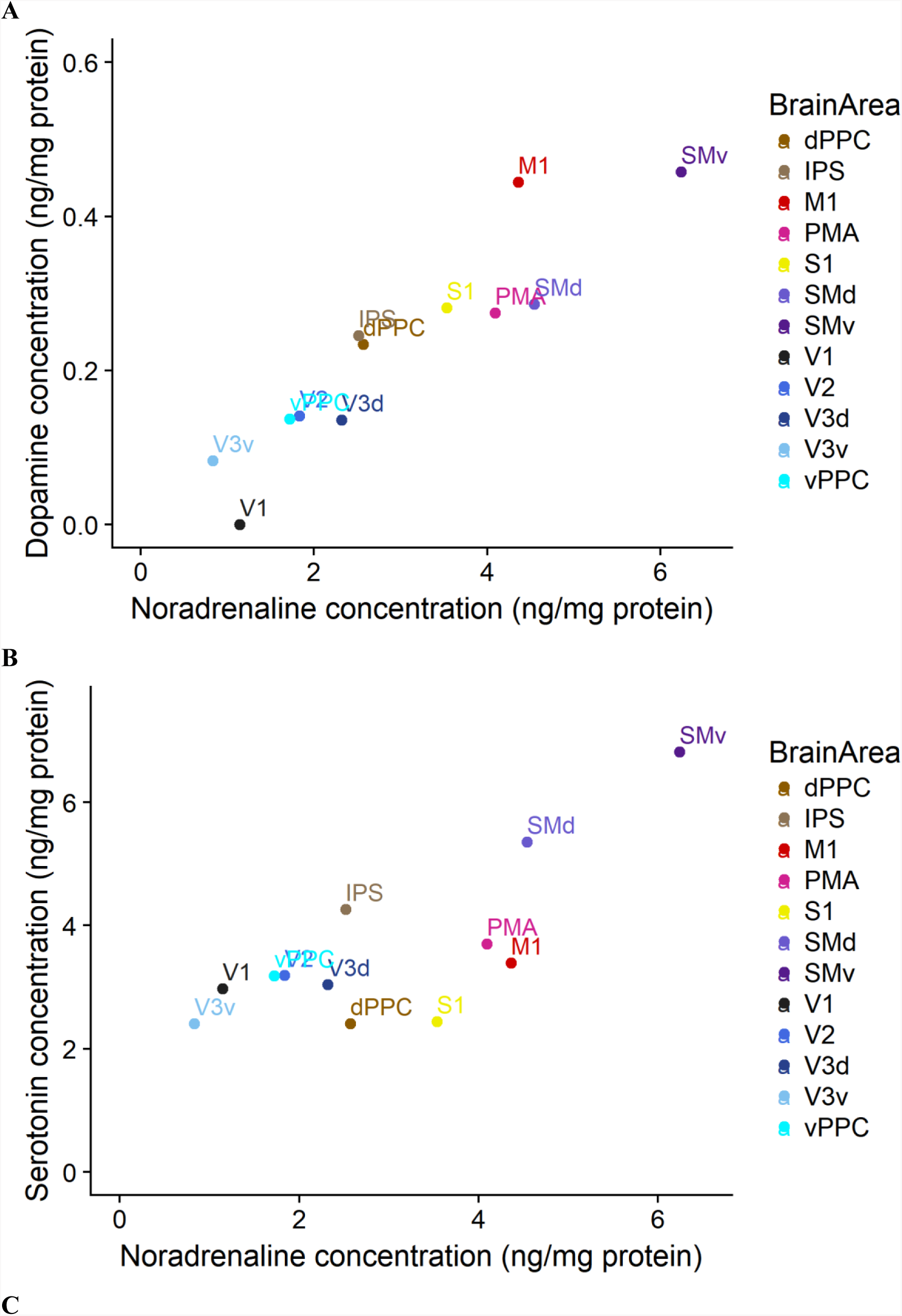
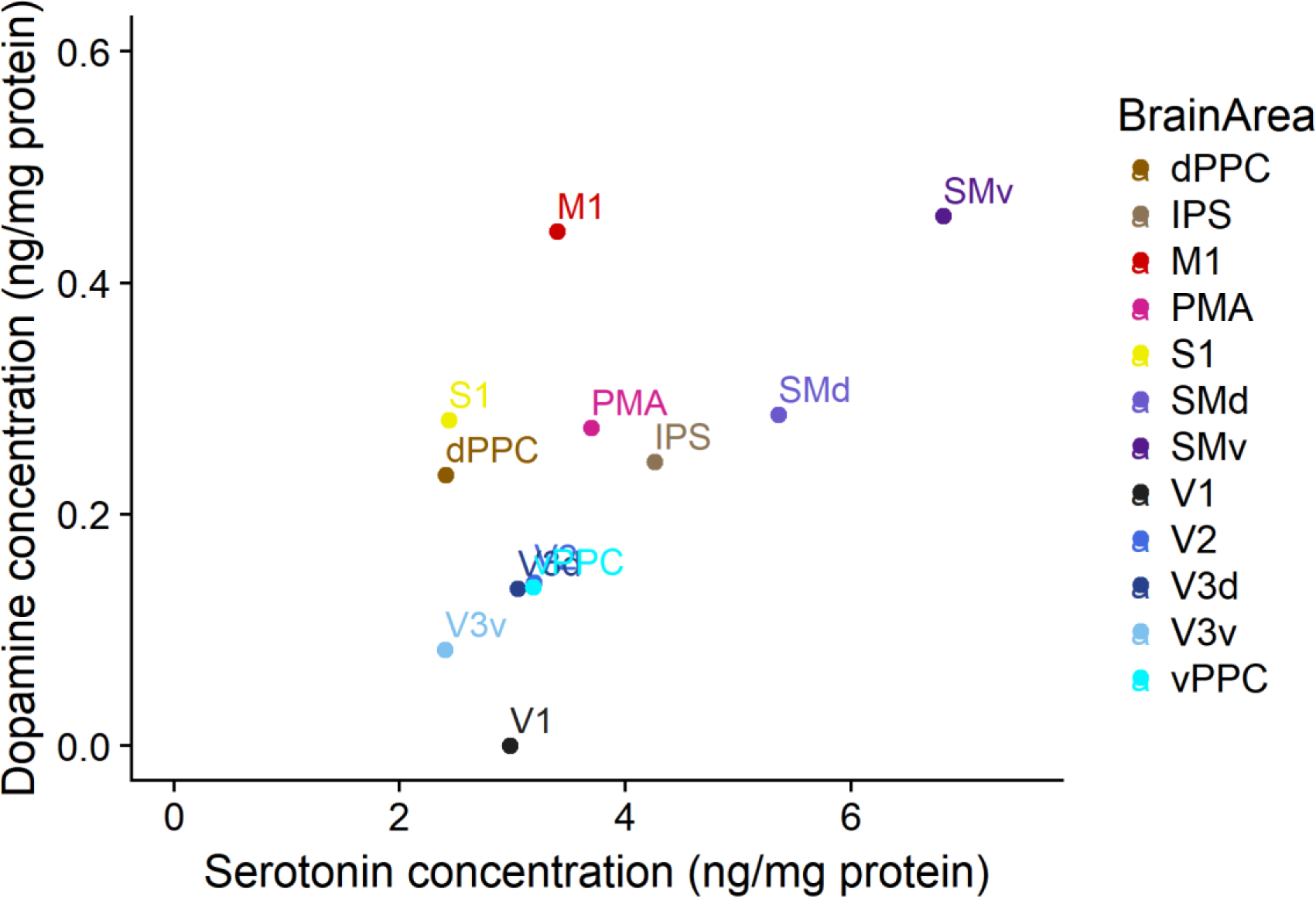
Plots showing center of mass data derived from monoamine concentrations. Colored dots (legend) designate center of mass of concentrations for a given brain area. Plots show comparisons of (**A**) dopamine and noradrenaline, (**B**) serotonin and noradrenaline, and (**C**) dopamine and serotonin.

**Figure 5.**
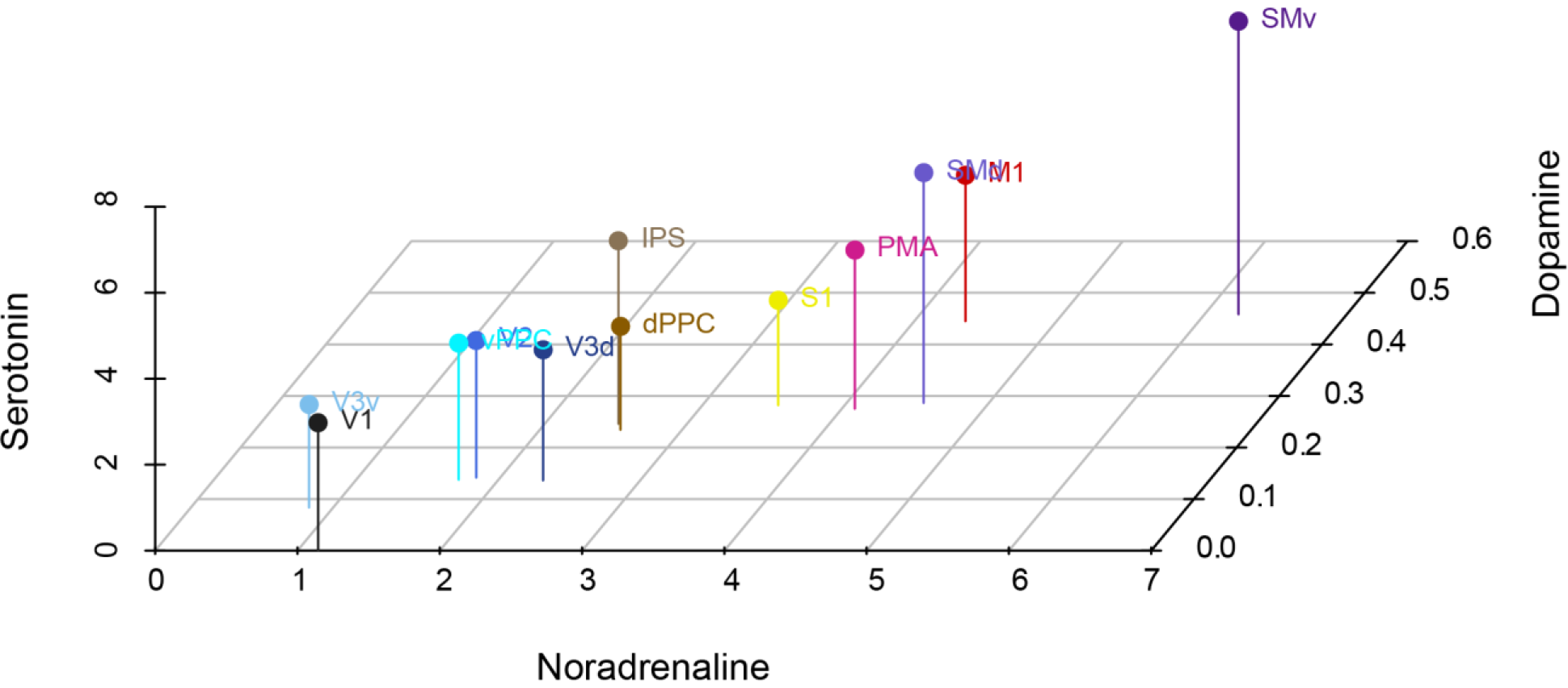
Center of mass concentration (in ng/mg protein) data represented in a multi-modulator space (x: noradrenaline; y: serotonin; z: dopamine). Colored dots designate center of mass of concentrations for a given brain area.

In semiquantitative measures of dopaminergic innervation density, Berger et al. (1988) noted denser dopaminergic innervation of agranular compared to granular cortical areas, which they stated does not parallel previous macaque neurochemical data (Brown et al. 1979, summarized in **Table 3**). In particular, Brown et al. (1979) noted higher levels of dopamine in granular prefrontal areas than in motor areas, while the opposite was true in the Berger et al. (1988) innervation study. However, these innervation results match better with subsequent studies that found lower levels of dopamine in prefrontal cortex than in motor areas (Pifl et al. 1991) and frontal cortex samples containing motor areas (Javoy-Agid et al. 1989). The granular vs. agranular difference in dopaminergic innervation was also noted in a study of human cortex (Gaspar et al. 1989). When comparing these results with other innervation studies (Levitt et al. 1984, Lewis et al. 1987), it appears that there is general concordance of results across studies (**Table 3**). Our results match fairly well with these studies with the only exception being that we see similar dopamine levels between agranular PMA and granular S1 (**Figure 4**). PMA samples in our study came from one macaque (**Figure 1**, **Table 1**), so it is possible that our result could be different with addition of samples from more macaques. Across all studies, it appears that dopaminergic innervation is moderately predictive of dopamine tissue concentration.

**Table 3.**
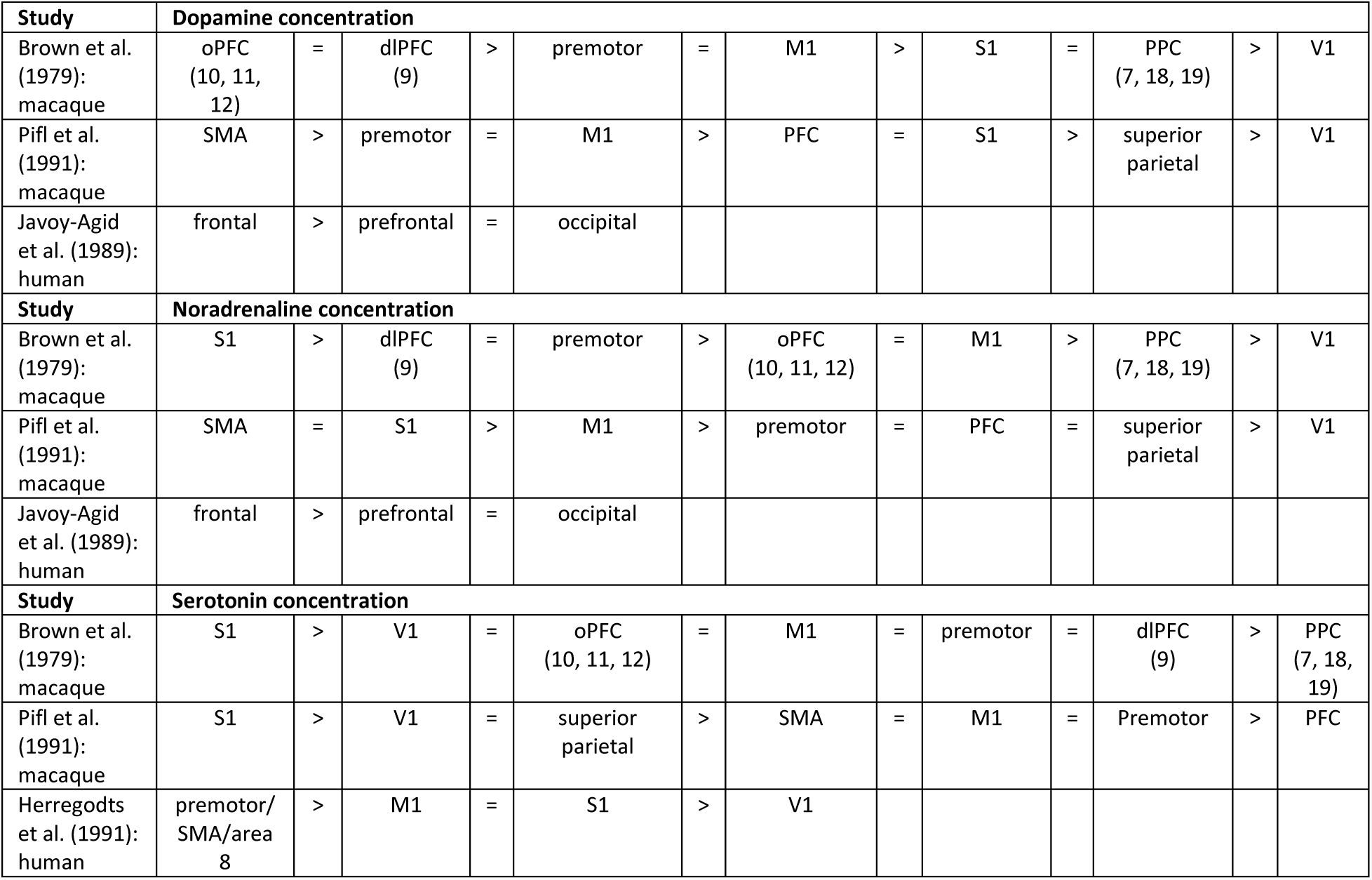
Summary of dopamine, noradrenaline, and serotonin concentrations from steady-state measure studies.

### Comparisons between serotonergic innervation and steady-state measures

In macaque studies of serotonergic innervation across multiple cortical areas, V1 exhibited the densest serotonergic innervation of all areas studied (Takeuchi and Sano 1983, Morrison and Foote 1986, Berger et al. 1988). Intriguingly, while Takeuchi and Sano (1983) and Berger et al. (1988) observed higher serotonergic fiber density in V1 than S1, in terms of serotonin concentration, the reverse was observed in previous neurochemical studies in macaque (Brown et al. 1988, Pifl et al. 1991). Likewise, a study in human found lower levels of serotonin in the visual cortex than S1 and motor areas (Herregodts et al. 1991). Similar to the innervation studies, our neurochemical data show higher levels of serotonin in V1 than in S1; however, unlike those studies, our data indicate that V1 has mid-range serotonin concentrations when compared to other studied regions (**Figure 4**).

Though no one serotonergic innervation study made clear comparisons across all areas detailed in **Table 2**, the studies do appear to make similar conclusions for the comparisons that were made. By comparison, steady state measures of concentration were much more variable (**Table 3**). Our own results show lower levels of serotonin than would be predicted for V1 and S1 based on innervation (**Figure 4**, **Table 2**). The difference in methodology between histological innervation and steady-state concentration studies may underlie this variability. With fresh tissue collection, it is difficult to ensure that comparable tissue has been collected in every instance, especially with an area like S1, within which somatosensory areas 1-3 exist. Likewise, because fresh tissue is subject to degradation, we are to some extent measuring the results of monoamine turnover. From this study and other steady-state studies, it is not clear that serotonergic innervation can be used to predict tissue concentrations of serotonin.

### Comparisons between noradrenergic innervation and steady-state measures

Studies of the macaque noradrenergic system and its innervation of cortex have typically been limited to specific regions (primary auditory: Campbell et al. 1987; V1: Foote and Morrison 1984, Kosofsky et al. 1984; prefrontal: Lewis and Morrison 1989). The most relevant standalone study of noradrenergic innervation across many cortical areas was conducted in humans (Gaspar et al. 1989) and is used here as a reference point for interpreting our results. In that study, noradrenergic innervation was highest in somatosensory and motor areas, and decreased in both rostral and caudal directions, with prefrontal and occipital areas demonstrating some of the lowest levels of innervation. This pattern was partially supported by comparisons to macaque data first presented in a book chapter by Lewis and Sesack (1997), which showed sparse noradrenergic innervation of prefrontal areas 9 and 10, and comparably denser innervation of M1 and S1.

Our data demonstrate that sensorimotor areas in and around the central sulcus have the highest levels of noradrenaline of the areas we sampled (**Figure 4**). Occipital areas exhibit some of the lowest noradrenaline levels measured in our samples (**Figure 4**). Both Brown et al. (1979) and Pifl et al. (1991) also measured their lowest noradrenaline concentrations in occipital samples. We did not examine prefrontal samples, so we cannot evaluate whether noradrenaline concentration follows a decreasing trend when moving away from the sensorimotor areas toward the rostral pole. In Brown et al. (1979), samples collected from both orbital and dorsolateral prefrontal cortex exhibited similar noradrenaline concentrations when compared to M1 and PMA samples. In another macaque study, Pifl et al. (1991) made a similar observation: noradrenaline levels in prefrontal samples were comparable to those in both motor and premotor samples. While innervation of prefrontal cortex appears less dense than that of sensorimotor areas in both human (Gaspar et al. 1989) and macaque (Lewis and Sesack 1997), noradrenaline concentration measures from macaque prefrontal cortex (Brown et al. 1979, Pifl et al. 1991) do not reflect the same stark rostral decrease observed in one study from humans (Javoy-Agid et al. 1989). Altogether, our noradrenergic concentration data match well with other steady-state studies of concentration (**Table 3**) as well as with innervation studies (**Table 2**). This indicates that density of noradrenergic innervation may help predict tissue concentration of noradrenaline.

### Comparisons with receptor localization and availability

While monoaminergic innervation patterns and concentrations have not been explored in V3d and V3v, autoradiographic study of the human visual cortex has explored receptor distribution and availability of the receiving circuits in these areas. For noradrenaline α1 and α2 as well as serotonin 5-HT1a receptors, the mean receptor availability does not differ between V3d and V3v; however, the laminar binding pattern for all three receptors does differ significantly between these two regions (Eickhoff et al. 2008). In the present study, we plotted concentrations of noradrenaline and serotonin for these regions (**Figure 3B**) and were able to observe how these regions compare to each other by examining center-of-mass values for samples (**Figure 4B**). V3d and V3v are separated from each other in these plots, with V3v potentially grouping with V1, and V3d more noticeably grouping with V2 and the posterior parietal cortex (**Figure 4B**). In our clustering analysis, this proposed grouping indeed arises (**Figure 6**). One explanation of these results is that these dorsal-ventral differences in V3 for both receptor distribution and neurochemical concentration indicate that V3d and V3v represent distinct noradrenergic and serotonergic signaling compartments. Another explanation is that these regions are not overtly distinct neurochemical environments, and we have failed to isolate samples containing only V3 tissue. As reported by Gattass et al. (1988), macaque V3 occupies a thin strip of cortex approximately 4-5 mm wide and is flanked posteriorly by V2 and anteriorly by several extrastriate areas. In particular, macaque V3d exhibits notable interanimal variability in its width, such that extrastriate areas that flank V3d anteriorly may sometimes occupy cortical space often devoted to V3d (Gattass et al. 1988). Our tissue collection strategy relied on morphological features rather than functional identification to parcellate cortical areas. It is possible that some of the separation noted between V3d and V3v can be attributed to samples containing tissue belonging to neighboring extrastriate tissue rather than purely V3 tissue. Caution should thus be exercised in attempting to understand these data in terms of the competing hypotheses surrounding the definition of V3 as an area (Lyon and Connolly 2012).

**Figure 6.**
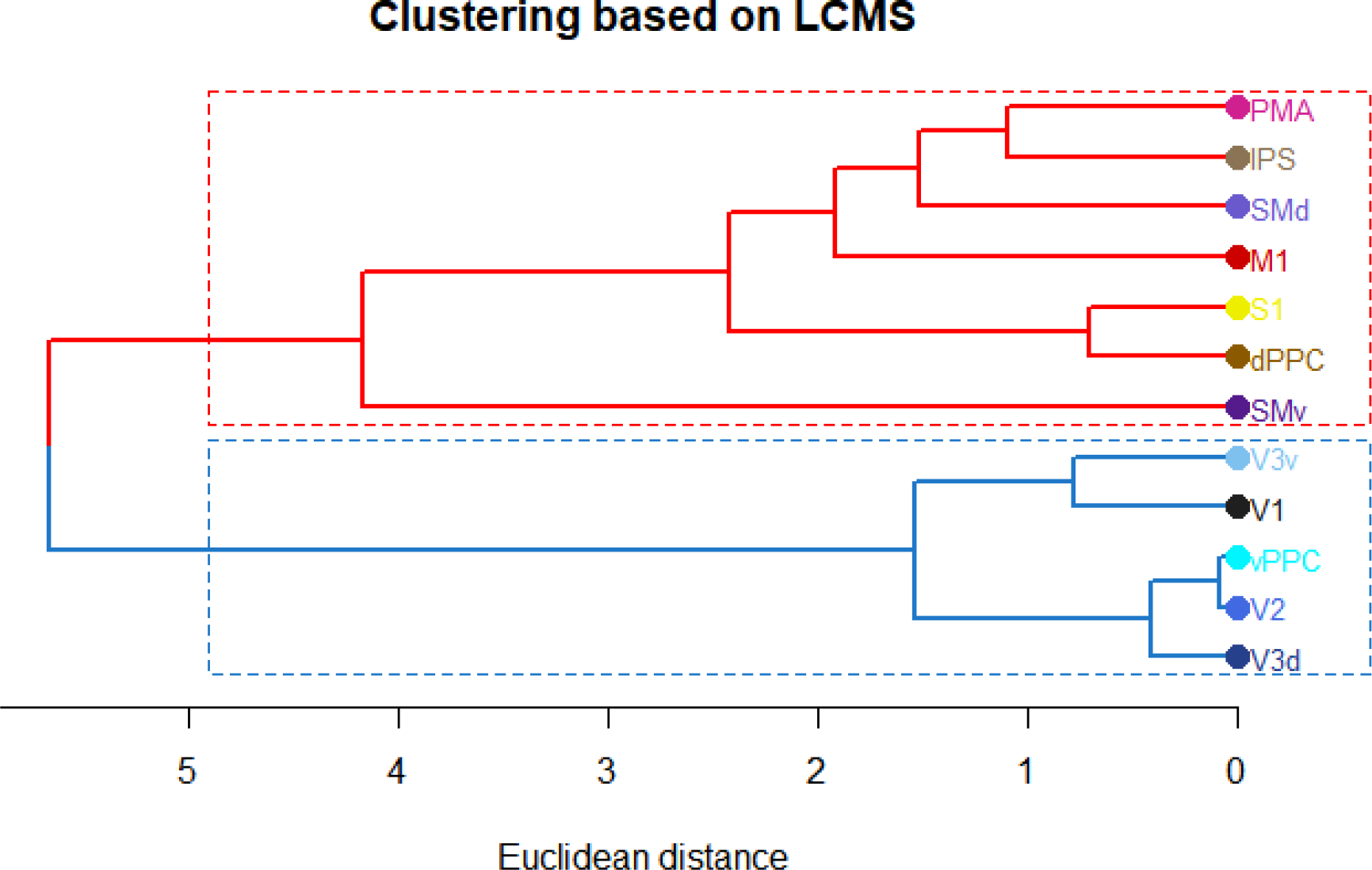
Hierarchical clustering of cortical areas by monoamine concentrations. Dendrogram shows cortical areas on right with lines extending from them to points at which these cortical areas are merged with another area based on similarity. Scale indicates the Euclidean distances (a measure of dissimilarity between areas) at which these merging points occur. Merging points that occur at lower distance values indicate areas that are more similar than those with merging points at higher distance values. Colored lines and boxes indicate clusters of areas that show similarity to each other based on measured monoamine concentrations.

Outside of the occipital cortex, receptor availability also differs between cortical areas. In a receptor autoradiography study of the sensorimotor areas, 5-HT2 serotonergic receptor binding availability was lower in M1 compared to S1 in both macaques and humans (Zilles et al. 1995). In our macaque samples, serotonin levels appear to be higher in M1 than S1 (**Figure 4**). This may indicate a mismatch between receptor density and concentration of serotonin. This mismatch was not seen in neurochemical studies by Brown et al. (1979) or Pifl et al. (1991), which more closely follow the receptor density described by Zilles et al. (1995). It is possible that there is enough variability within S1 regions that we sampled from relatively low concentration areas. In a study of human S1, 5-HT2 receptor availability was lower laterally than medially in all four areas examined: 1, 2, 3a, and 3b (Geyer et al. 1997). Our study examined monoamine concentrations of these sensorimotor areas in ventrolateral (SMv) and dorsomedial (SMd) samples. Our data indicate that the lateral samples exhibit a higher concentration of serotonin compared to dorsal samples. Again, this indicates that we observed a mismatch between our serotonin concentration data and the receptor densities noted by Geyer et al. (1997). In that study, 5-HT2 receptor availability varied between the sensorimotor areas, and within area 1 density was higher in the posterior portion than the anterior portion. This indicates that within and across these sensory areas, serotonin receptor densities vary.

Unlike the Geyer et al. (1997) study, our collection process (and those of Brown et al. 1979 and Pifl et al. 1991) did not allow us to distinguish between sensorimotor areas in order to make any fine-grained comparisons within these areas. A likely possibility is that due to these methods, our observations for SMv and SMd blur across several regions that are distinct monoaminergic signaling compartments. This is an important caveat for us to consider in terms of our parietal cortex results as well. Within the inferior parietal lobule, mean 5-HT1a receptor density varies across its rostral-to-caudal extent: density is low in the rostral, intermediate in the middle, and high in the caudal sector (Geyer et al. 2005). Samples collected from vPPC in our study may also represent a combination of these varied signaling environments.

Cluster analyses can be applied to receptor autoradiography data in order to understand similarities and dissimilarities in receptive circuitry between brain areas (Scheperjans et al 2005, Caspers et al 2013). We applied similar analyses to our neurochemical data to compare it with clustering based on receptor autoradiography. Our clustering data suggest one cluster that contains visual areas (V1, V2, V3v, V3d) and PPCv, and another cluster that contains sensorimotor areas (M1, S1, SMv, SMd, PMA) as well as PPCd and IPS (**Figure 6**). Using receptor autoradiography-based clustering, Scheperjans et al. (2005) noted clustering between human somatosensory and superior parietal area 5 samples. Clusters containing primary and extrastriate visual areas tended to share more similarities with each other than with somatosensory or parietal samples. In a more recent human study, Caspers et al. (2013) also clustered autoradiography data and noted two large clusters: one contained multiple sensorimotor, auditory, and visual areas, while the other contained areas from the superior and inferior parietal area as well as V3v.

### Functional implications and future considerations

In evaluating the data from our study, it is important to consider the nature of the conclusions we can readily make. Our data are derived from postmortem tissue and therefore represent a steady-state *ex vivo* rather than a dynamical *in vivo* measurement. Additionally, our monoamine concentration values represent molecules found both inside monoaminergic axons innervating cortex and outside in the extracellular signaling space. Therefore, we cannot make conclusions about how these monoamines are dynamically signaling in an extracellular space.

Our data can be used is to improve interpretation of functional studies. For example, Marrocco et al. (1987) demonstrated catecholamine release in macaque V1 but not V2 in response to visual stimuli. Changes in catecholamines were monitored using electrochemical recordings that cannot differentiate between dopamine and noradrenaline (Fox and Wightman 2017). Our results indicate that catecholamine changes measured in V1 by Marrocco et al. (1987) can likely be attributed to noradrenaline rather than dopamine. For reference, electrochemical recording studies in brain areas with a high noradrenaline to dopamine ratio (approximately 10:1 or greater) attribute electrochemical changes to noradrenaline release (Park et al. 2009). Based on our results (**Figure 4A**), this attribution of signal change to noradrenaline is likely fair for V1. Additionally, the lack of measured change in the V2 catecholamine signal is likely due to noradrenaline release not changing. Our results indicate that V2 exhibits similar, if not higher, levels of noradrenaline when compared to V1 and a favorable noradrenaline to dopamine ratio (**Figure 4A**). When these data are combined with evidence that V2 exhibits denser noradrenergic innervation than V1, it seems unlikely that Marrocco et al. (1987) would have experienced difficulty measuring a noradrenergic signal change in V2 had it existed. In this way, our data point to the utility of connecting macaque neurochemical characterization to studies of cortical function.

## Acknowledgments

We thank Troy Hackett and Ramnarayan Ramachandran for provision of tissue, Pooja Balaram for tissue slab acquisition, Rachel Gilfarb for assistance with tissue punch collection and data analysis, Ginger Milne and Benlian Gao of the Vanderbilt Neurochemistry Core (supported in part by the EKS NICHD of NIH under award #U54HD083211) for assistance with biogenic amine quantification, and Mary Feurtado for assistance in animal care. This work was supported by NIH grant MH093567 (to A. Disney).

## References

Berger, B., Trottier, S., Verney, C., Gaspar, P., & Alvarez, C. (1988). Regional and laminar distribution of the dopamine and serotonin innervation in the macaque cerebral cortex: a radioautographic study. Journal of Comparative Neurology, 273(1), 99–119.

Björklund, A., Divac, I., & Lindvall, O. (1978). Regional distribution of catecholamines in monkey cerebral cortex, evidence for a dopaminergic innervation of the primate prefrontal cortex. Neuroscience letters, 7(2), 115–119.

Brown, R. M., Crane, A. M., & Goldman, P. S. (1979). Regional distribution of monoamines in the cerebral cortex and subcortical structures of the rhesus monkey: concentrations and in vivo synthesis rates. Brain research, 168(1), 133–150.

Campbell, M. J., Lewis, D. A., Foote, S. L., & Morrison, J. H. (1987). Distribution of choline acetyltransferase-, serotonin-, dopamine-β-hydroxylase-, tyrosine hydroxylase-immunoreactive fibers in monkey primary auditory cortex. Journal of Comparative Neurology, 261(2), 209–220.

Caspers, S., Schleicher, A., Bacha-Trams, M., Palomero-Gallagher, N., Amunts, K., & Zilles, K. (2012). Organization of the human inferior parietal lobule based on receptor architectonics. Cerebral Cortex, 23(3), 615–628.

Coppola, J. J., Ward, N. J., Jadi, M. P., & Disney, A. A. (2016). Modulatory compartments in cortex and local regulation of cholinergic tone. Journal of Physiology-Paris, 110(1), 3–9.

Dylla, M., Hrnicek, A., Rice, C., & Ramachandran, R. (2013). Detection of tones and their modification by noise in nonhuman primates. Journal of the Association for Research in Otolaryngology, 14(4), 547–560.

Eickhoff, S. B., Rottschy, C., Kujovic, M., Palomero-Gallagher, N., & Zilles, K. (2008). Organizational principles of human visual cortex revealed by receptor mapping. Cerebral Cortex, 18(11), 2637–2645.

Foote, S. L., & Morrison, J. H. (1984). Postnatal development of laminar innervation patterns by monoaminergic fibers in monkey (Macaca fascicularis) primary visual cortex. Journal of Neuroscience, 4(11), 2667–2680.

Fox, M. E., & Wightman, R. M. (2017). Contrasting Regulation of Catecholamine Neurotransmission in the Behaving Brain: Pharmacological Insights from an Electrochemical Perspective. Pharmacological reviews, 69(1), 12–32.

Freedman, R., Foote, S. L., & Bloom, F. E. (1975). Histochemical characterization of a neocortical projection of the nucleus locus coeruleus in the squirrel monkey. Journal of comparative neurology, 164(2), 209–231.

Fuxe, K., & Agnati, L. F. (Eds.). (1991). Volume transmission in the brain: novel mechanisms for neural transmission (Vol. 1). Raven Press.

Galili, T. (2015). dendextend: an R package for visualizing, adjusting and comparing trees of hierarchical clustering. Bioinformatics, 31(22), 3718–3720.

Gaspar, P., Berger, B., Febvret, A., Vigny, A., & Henry, J. P. (1989). Catecholamine innervation of the human cerebral cortex as revealed by comparative immunohistochemistry of tyrosine hydroxylase and dopamine-beta- hydroxylase. Journal of Comparative Neurology, 279(2), 249–271.

Gattass, R., Sousa, A. P., & Gross, C. G. (1988). Visuotopic organization and extent of V3 and V4 of the macaque. Journal of Neuroscience, 8(6), 1831–1845.

Gatter, K. C., & Powell, T. P. S. (1977). The projection of the locus coeruleus upon the neocortex in the macaque monkey. Neuroscience, 2(3), 441–445.

Geyer, S., Luppino, G., Ekamp, H., & Zilles, K. (2005). The macaque inferior parietal lobule: cytoarchitecture and distribution pattern of serotonin 5-HT1A binding sites. Anatomy and embryology, 210(5-6), 353–362.

Herregodts, P., Ebinger, G., & Michotte, Y. (1991). Distribution of monoamines in human brain: evidence for neurochemical heterogeneity in subcortical as well as in cortical areas. Brain research, 542(2), 300–306.

Javoy-Agid, F., Scatton, B., Ruberg, M., L’heureux, R., Cervera, P., Raisman, R., Maloteaux, J.-M., Beck, H., & Agid, Y. (1989). Distribution of monoaminergic, cholinergic, and GABAergic markers in the human cerebral cortex. Neuroscience, 29(2), 251–259.

Jones, B. E., & Moore, R. Y. (1977). Ascending projections of the locus coeruleus in the rat. II. Autoradiographic study. Brain research, 127(1), 23–53.

Kosofsky, B. E., Molliver, M. E., Morrison, J. H., & Foote, S. L. (1984). The serotonin and norepinephrine innervation of primary visual cortex in the cynomolgus monkey (Macaca fascicularis). Journal of Comparative Neurology, 230(2), 168–178.

Levitt, P., Rakic, P., & Goldman- Rakic, P. (1984). Region-specific distribution of catecholamine afferents in primate cerebral cortex: A fluorescence histochemical analysis. Journal of Comparative Neurology, 227(1), 23–36.

Lewis, D. A., Campbell, M. J., Foote, S. L., Goldstein, M., & Morrison, J. H. (1987). The distribution of tyrosine hydroxylase-immunoreactive fibers in primate neocortex is widespread but regionally specific. Journal of Neuroscience, 7(1), 279–290.

Lewis, D. A., & Morrison, J. H. (1989). Noradrenergic innervation of monkey prefrontal cortex: A dopamine-β-hydroxylase immunohistochemical study. Journal of Comparative Neurology, 282(3), 317–330.

Lewis, D. A., & Sesack, S. R. (1997). Chapter VI dopamine systems in the primate brain. Handbook of chemical neuroanatomy, 13, 263–375.

Ligges, U. and Mächler, M. (2003). Scatterplot3d - an R Package for Visualizing Multivariate Data. Journal of Statistical Software 8(11), 1–20.

Lyon, D. C., & Connolly, J. D. (2012). The case for primate V3. Proceedings of the Royal Society of London B: Biological Sciences, 279(1729), 625–633.

Marrocco, R. T., Lane, R. F., McClurkin, J. W., Blaha, C. D., & Alkire, M. F. (1987). Release of cortical catecholamines by visual stimulation requires activity in thalamocortical afferents of monkey and cat. Journal of Neuroscience, 7(9), 2756–2767.

Morrison, J. H., & Foote, S. L. (1986). Noradrenergic and serotoninergic innervation of cortical, thalamic, and tectal visual structures in Old and New World monkeys. Journal of Comparative Neurology, 243(1), 117–138.

Murtagh, F., & Legendre, P. (2014). Ward’s hierarchical agglomerative clustering method: which algorithms implement Ward’s criterion?. Journal of Classification, 31(3), 274–295.

Park, J., Kile, B. M., & Mark Wightman, R. (2009). In vivo voltammetric monitoring of norepinephrine release in the rat ventral bed nucleus of the stria terminalis and anteroventral thalamic nucleus. European Journal of Neuroscience, 30(11), 2121–2133.

Phillis, J. W. (1968). Acetylcholine release from the cerebral cortex: its role in cortical arousal. Brain research, 7(3), 378–389.

Pifl, C., Schingnitz, G., & Hornykiewicz, O. (1991). Effect of 1-methyl-4-phenyl-1, 2, 3, 6-tetrahydropyridine on the regional distribution of brain monoamines in the rhesus monkey. Neuroscience, 44(3), 591–605.

R Core Team (2017) R: A language and environment for statistical computing. Vienna: R Foundation for Statistical Computing.

Romanski, L. M., Averbeck, B. B., & Diltz, M. (2005). Neural representation of vocalizations in the primate ventrolateral prefrontal cortex. Journal of Neurophysiology, 93(2), 734–747.

Rousseeuw, P. J. (1987). Silhouettes: a graphical aid to the interpretation and validation of cluster analysis. Journal of computational and applied mathematics, 20, 53–65.

Saper, C. B. (1987). Diffuse cortical projection systems: anatomical organization and role in cortical function. In Handbook of Physiology. The Nervous System V (Plum, F., ed.), pp. 169–210, American Physiology Society.

Scheperjans, F., Palomero-Gallagher, N., Grefkes, C., Schleicher, A., & Zilles, K. (2005). Transmitter receptors reveal segregation of cortical areas in the human superior parietal cortex: relations to visual and somatosensory regions. Neuroimage, 28(2), 362–379.

Takeuchi, Y., & Sano, Y. (1983). Immunohistochemical demonstration of serotonin nerve fibers in the neocortex of the monkey (Macaca fuscata). Anatomy and embryology, 166(2), 155–168.

Schultz, W. (2002). Getting formal with dopamine and reward. Neuron, 36(2), 241–263.

Ward Jr, J. H. (1963). Hierarchical grouping to optimize an objective function. Journal of the American statistical association, 58(301), 236–244.

Wickham H (2009) ggplot2: elegant graphics for data analysis. New York: Springer.

Wilke, C.O. (2016). Cowplot: Streamlined plot theme and plot annotations for ‘ggplot2’. http://cran.r-project.org/web/packages/cowplot/index.html.

Wilson, M. A., & Molliver, M. E. (1991). The organization of serotonergic projections to cerebral cortex in primates: retrograde transport studies. Neuroscience, 44(3), 555–570.

Zilles, K., Schlaug, G., Matelli, M., Luppino, G., Schleicher, A., Qü, M., Dabringhaus, A., Seitz, R., & Roland, P. E. (1995). Mapping of human and macaque sensorimotor areas by integrating architectonic, transmitter receptor, MRI and PET data. Journal of Anatomy, 187(Pt 3), 515.

